# Efficacy of chronic 5-HT_1A_ receptor agonism by NLX-112 in a mouse model of Spinocerebellar Ataxia type 3

**DOI:** 10.1101/2025.08.21.671624

**Authors:** B Ferreira-Lomba, S Guerreiro, S Duarte-Silva, D Cunha-Garcia, S Oliveira, C Vieira, J Pereira-Sousa, D Vilasboas-Campos, A Vidinha-Mira, D Monteiro-Fernandes, MA Varney, MS Kleven, A Newman-Tancredi, A Teixeira-Castro, P Maciel

## Abstract

**Background:** Spinocerebellar ataxia type 3 (SCA3) is an autosomal dominant neurodegenerative disorder caused by an elongated polyglutamine (polyQ) sequence in the ataxin-3 protein. This expansion triggers neuropathological events, leading to progressive motor disturbances. Currently, no approved therapy exists for this debilitating condition, but compelling evidence suggests that targeting the serotonergic system can significantly attenuate SCA3 disease progression in animal models.

**Objective:** This study aimed to assess the effects of NLX-112, a highly selective serotonin 1A receptor (5-HT_1A_R) full agonist, in the CMVMJD135 transgenic mouse model of SCA3.

**Methods:** NLX-112 (0.625 and 5 mg/kg/day) and tandospirone (a 5-HT_1A_R partial agonist used as a comparator; 20 and 80 mg/kg/day) were administered chronically in drinking water for 34 weeks, starting prior to symptom onset. To evaluate the effects of the drugs on SCA3 mice, motor-related behavioral tests and neuropathological techniques were employed.

**Results:** Treatment with the higher dose of NLX-112 led to improvements in motor coordination and balance, and slowing of symptom deterioration as the disease progressed. These beneficial effects were not achieved with tandospirone. NLX-112 treatment also elicited neuroprotective effects, reducing dopaminergic (tyrosine hydroxylase-positive) cell loss and astrocyte reactivity in the substantia nigra.

**Conclusions:** NLX-112 treatment, started pre-symptomatically, enhanced motor function, slowed disease progression and elicited neuroprotective effects in SCA3 mice, supporting its further development as a drug candidate for treatment of ataxia and related movement disorders.

**Graphical Abstract:** 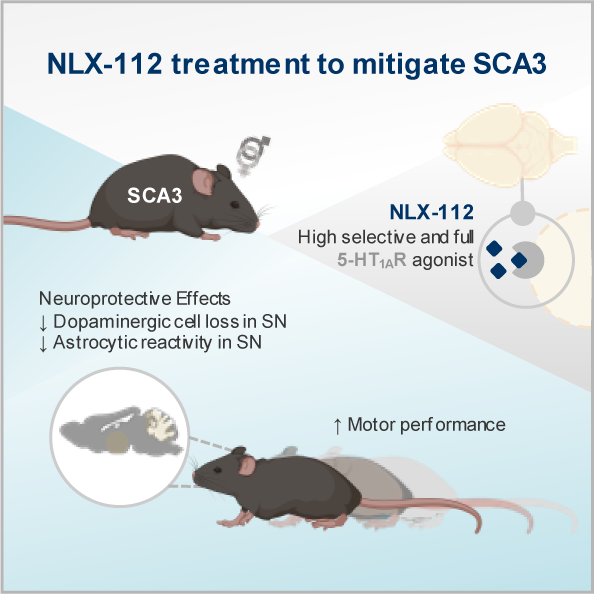

*Key findings:*

- NLX-112 attenuated motor deficits of SCA3 mice, when administered chronically prior to disease onset.
- NLX-112 reduced neuropathological biomarkers in SCA3 mice, namely by restoring dopaminergic neuron loss and decreasing astrocyte reactivity.
- NLX-112 is a potential candidate for addressing ataxia-related deficits in SCA3 patients.

## Introduction

Spinocerebellar ataxia type 3 (SCA3) is the most common form of dominantly inherited ataxia (1–3). This neurodegenerative disease is caused by an unstable expansion of a cytosine-adenine-guanidine (CAG) trinucleotide repeat within the coding region of the ataxin-3 (*ATXN3*) gene (4, 5), which is translated into an abnormal ATXN3 protein carrying an extended polyglutamine (polyQ) tract (6). The CAG repeat size ranges from 10 to 44 in healthy individuals, whereas in SCA3 patients it varies from 61 to 87 (5), and its length is inversely correlated with the age at disease onset (7). A core feature of SCA3 is progressive ataxia, characterized by motor incoordination, affecting gait, balance, fine movements and speech; rigidity, spasticity, dystonia and loss of strength are often present (1). ATXN3 is a deubiquitylating enzyme (DUB) (8), involved in multiple cellular functions, including protein degradation, responses to heat and oxidative stresses, DNA damage, transcription, cytoskeletal organization and cell adhesion (9, 10). The ATXN3 polyQ expansion causes conformational alterations, increasing its propensity to aggregate and accumulate within the nucleus and axonal tracts of neurons (11). This results in a toxic gain-of-function thought to be responsible for neuronal dysfunction and to mediate neurodegeneration within the central nervous system (9, 12), including the deep cerebellar nuclei (DCN), pontine nuclei (PN), *substantia nigra* (SN) and spinocerebellar tracts. The underlying molecular and cellular mechanisms are still unclear, hampering the path towards finding effective therapies. Currently, only physical and speech therapies or medications for dystonia and extrapyramidal symptoms are available (13). Disease-modifying treatments are urgently needed for SCA3 patients.

We recently found that the selective serotonin reuptake inhibitor (SSRI) citalopram, which inhibits serotonin (5-HT) transporter (SERT) function to increase 5-HT synaptic availability, reduced mutant ATXN3 aggregation and improved motor function in *C. elegans* and mouse models of SCA3 (14, 15). The beneficial effect in *C. elegans* was dependent on MOD-5, the SERT homologue, and on the 5-HT receptor SER-4, the *C. elegans* 5-HT_1A_ receptor (5-HT_1A_R) homologue (14).

We therefore focused the present study on the 5-HT_1A_R, the most abundant 5-HT receptor subtype in the brain (16). Presynaptic 5-HT_1A_ autoreceptors are found on the cell bodies and dendrites of raphe nuclei 5-HT neurons, while postsynaptic 5-HT_1A_ heteroreceptors are expressed in target areas receiving 5-HT innervation, such as the hippocampus and prefrontal cortex (17). 5-HT_1A_ autoreceptors regulate 5-HT neuron firing, while the heteroreceptors mediate neuroplasticity, fine motor control and coordination (18, 19). These receptors also play a role in thermoregulation, with selective 5-HT_1A_R agonists inducing hypothermia (20).

5-HT_1A_Rs have attracted interest as a promising therapeutic target for movement disorders. Previous small open-label studies using tandospirone, a partial 5-HT_1A_R agonist, showed promising results in SCA3 and spinocerebellar ataxia type 6 (SCA6) patients, reducing cerebellar ataxia and other disease-related symptomatology (20, 21). However, tandospirone has only modest receptor specificity, interacting with different receptors (22), which may limit its efficacy and lead to undesirable side effects. In contrast, NLX-112 (befiradol, F-13640), an exceptionally selective agonist with high affinity for 5-HT_1A_Rs (23, 24) has been developed. NLX-112 showed its efficacy in multiple models of movement disorders, such as Parkinson’s disease (PD), in which it reversed haloperidol-induced dystonia and L-DOPA-induced dyskinesia in parkinsonian rats (25) and revealed anti-dyskinetic and anti-parkinsonian-like properties in 1-methyl-4-phenyl-1,2,3,6-tetrahydropyridine (MPTP)-treated marmosets and macaques (25–28). Additionally, clinical trials with NLX-112 indicated that it is safe and well-tolerated (29, 30), and significantly reduced PD patients’ dyskinesias and parkinsonism (31, 32).

As concerns ataxia, in a *C. elegans* model of SCA3, acute and chronic treatment with NLX-112 rescued motor function and suppressed mutant ATXN3 aggregation, an effect dependent on the presence 5-HT_1A_Rs (33). Here, we used the well-characterized CMVMJD135 mouse model of SCA3 (14, 34–37) to evaluate whether early and sustained treatment with NLX-112 could improve motor dysfunction and neuropathology. After establishing an appropriate dosing strategy, we observed that NLX-112 (and not tandospirone) slowed motor deterioration in a dose-dependent manner, reduced tyrosine hydroxylase-positive (TH^+^) cells loss and astrocyte reactivity in the SN, suggesting a neuroprotective effect in SCA3 mice. These findings highlight its potential to treat SCA3 and eventually other movement disorders.

## Methods

### Study Design

This study aimed to assess the therapeutic efficacy of NLX-112 administration in the CMVMJD135 mouse model of SCA3, using tandospirone as a reference drug. At five weeks of age, mice were randomly assigned to treatment groups. A chronic dosing protocol was first optimized by assessing brain and plasma levels following acute oral gavage (OG), which informed the doses used in drinking water (DW) for long-term treatment. NLX-112 and tandospirone were then chronically administered via DW starting before the onset of motor symptoms. Motor behavior and welfare were evaluated at defined timepoints, and a subset of NLX-112-treated and untreated animals was used for blinded neuropathological analysis. All procedures followed FELASA guidelines, adhered to the 3Rs principles, and were approved by the ORBEA-ICVS ethics committee (ref. ORBEA EM/ICVS-I3Bs_007/2019) and the national authority DGAV (ref. 2022-02-07 003455). All researchers and facilities were DGAV-certified.

### Experimental Groups and General Methods

CMVMJD135 mice and wild-type (WT) littermates (genetic background strain C57BL/6J) of both sexes were used in this study. The experimental groups are presented in **Table 1**. DNA extraction, genotyping and determination of the number of CAG repeats were performed as previously reported (34, 38). The mice were generated in-house and by *in vitro* fertilization (at Charles River Laboratories, France).

**Table 1.** Description of experimental groups per experiment.

| Experiment | Number of animals/<br>experimental group | Experimental Groups |  | Treatment route | Average CAG repeats | Total N |
| --- | --- | --- | --- | --- | --- | --- |
|  |  | Genotype | Treatment/Dose |  |  |  |
| Drug Target Engagement | 12 ♀/♂ | SCA3 | Vehicle | Oral Gavage (OG) | 140 | 70 |
|  |  |  | NLX-112 0.625 mg/kg |  | 139 |  |
|  |  |  | NLX-112 2.5 mg/kg |  | 139 |  |
|  |  |  | NLX-112 10 mg/kg |  | 143 |  |
|  |  |  | TD 2.5 mg/kg |  | 139 |  |
|  |  |  | TD 20 mg/kg |  | 141 |  |
|  |  |  | TD 40 mg/kg |  | 138 |  |
|  | 12 ♀/♂ | WT | Vehicle |  | - |  |
| Chronic Treatment<br>(2-4 weeks) |  | SCA3 | Vehicle | Drinking Water (DW) | 141 | 60 and 21 |
|  |  |  | NLX-112 0.625 mg/kg |  | 141 |  |
|  |  |  | NLX-112 5 mg/kg |  | 142 |  |
|  |  |  | TD 2.5 mg/kg |  | 143 |  |
|  |  |  | TD 20 mg/kg |  | 143 |  |
|  |  |  | TD 80 mg/kg |  | 135 |  |
| Pre-symptomatic Treatment Initiation | 23-24 ♀/♂ | WT | Vehicle | Drinking Water (DW) | - | 167 |
|  |  | SCA3 | Vehicle 1 |  | 142 |  |
|  |  |  | NLX-112 0.625 mg/kg |  | 144 |  |
|  |  |  | NLX-112 5 mg/kg |  | 143 |  |
|  |  |  | Vehicle 2 |  | 144 |  |
|  |  |  | TD 20 mg/kg |  | 143 |  |
|  |  |  | TD 80 mg/kg |  | 146 |  |

Abbreviations: CAG, Cytosine-Adenine-Guanine; SCA3, Spinocerebellar Ataxia Type 3; TD, Tandospirone; ♀, Female; ♂, Male.

Housing and health conditions can be found in supplementary methods. At the end of each experiment, animals were euthanized for specific analyses. For drug exposure studies, mice were decapitated, and brain and plasma samples were flash-frozen in liquid nitrogen and stored at –80 °C. For neuropathology, animals were deeply anesthetized (150 mg/kg ketamine + 0.3 mg/kg medetomidine) and transcardially perfused with cold 0.9% saline followed by 4% paraformaldehyde (PFA) in PBS. Brains and spinal cords were collected, post-fixed in 4% PFA (72 h for brains, 2 weeks for SC), then cryoprotected in 30% sucrose with 0.02% sodium azide at 4 °C until sectioning.

### Compound Preparation and Administration

NLX-112 fumarate and tandospirone citrate were provided by Neurolixis. These compounds were used without further purification and dissolved in water when administered via OG and DW. The drugs’ dosages were adjusted to the body weight of each animal (for treatment via OG) or to the mean of the body weights of the animals housed in the same cage (for treatment via DW). For the treatment via OG, NLX-112 and tandospirone were prepared on the day of the experiment; for DW administration, the bottles with NLX-112 were substituted weekly and the bottles with tandospirone twice/week (the water intake/cage was measured at the moment of replacement).

### Drug Exposure Analysis

Brain and blood were collected for drug exposure studies. The blood was collected in EDTA-coated tubes, kept on ice for 1 hour and centrifuged at 13,000 rpm for 15 minutes: the obtained plasma was stored at -80°C. The quantification of the drug levels in the brain and plasma was outsourced to Eurofins ADME BIOANALYSES (France), using liquid chromatography-tandem mass spectrometry (LC-MS/MS) (detailed protocol can be found in supplementary methods).

### Body Weight and Temperature Measurement

Body weight was measured weekly. The body temperature of each animal was measured using a thermal rectal probe (Rodent Warmer X1, Stoelting Co.) covered in petroleum jelly, at predefined time points.

### Motor Function Evaluation

The motor function was evaluated by blinded experimenters, using the beam walking test, starting at week 7 and then performed biweekly from 10 weeks onwards until 40 weeks of age, as previously described (34). More details can be found in the supplementary methods. The number of animals used for behavioral assessment is reported in **Table 1**.

### Immunohistochemistry

Whole brains and SC of 41-week-old mice (mixed sexes) were sectioned in the sagittal and coronal planes, respectively, with a thickness of 40 μm, using the Vibrating Blade Microtome Leica VT 1000 S. The slices selected for immunohistochemistry (n= 5-6 animals, 5-10 slices/animal/experimental group) were incubated with Ultra Streptavidin HRP Kit (Multispecies DAB, Cat. 929501, BioLegend), following the protocol proposed by the manufacturer. Negative controls were used, consisting of tissue slices that followed the immunostaining protocol, without incubation with primary antibodies. The antibodies used are listed in **Table 2**. Upon immunolabeling, the slices were counterstained with hematoxylin 12.5-25 % (Leica AutoStainer XL ST5010), following a pre-established protocol. Microscopy and specific quantification software were used to perform all the neuropathological analyses (details on supplementary methods).

**Table 2.** Antibodies used for neuropathology analysis.

| Study | Antibody | Species | Dilution | Company/Product Reference |
| --- | --- | --- | --- | --- |
| Quantification of ATXN3-positive intranuclear inclusions | Anti-ATXN3 (1H9) | Mouse | 1:500 | Millipore, MAB5360 |
| Quantification of TH-positive neurons | Tyrosine hydroxylase (TH) | Rabbit | 1:300 | Merck, AB152 |
| Astrocyte reactivity | Glial Fibrillary Acidic Protein (GFAP) | Rabbit | 1:500 | DAKO, Z0334 |

### Statistical Analysis

The statistical analysis was performed using IBM SPSS Statistics 25 and the graphical representations were made using GraphPad Prism 10.4.1. (627). The sample size applied in each experiment was calculated utilizing G*Power 3.1.9.4, based on power analysis obtained from previous studies, and the effect size was determined intending an improvement of 50 %. The critical value used for significance was *p* ≤ 0.05. The statistical analyses for each experiment included both genotypes (SCA3 and WT), all the treatments (vehicle, NLX-112 and tandospirone) and respective doses. The normality of the data was analyzed using descriptive statistics (central tendency, dispersion, kurtosis and skewness) and Shapiro-Wilk Test (*p* > 0.05) and homogeneity of variances was measured by applying the Levene’s Test (*p* > 0.05). The outliers were removed when the interquartile ranges from the mean deviated more than 1.5. One-Way, Two-Way and Mixed design ANOVAs were performed to compare group measures, followed by Tukey’s or Sidak post-hoc tests. When the assumption of normal distribution was not met, the results were analyzed by Kruskal-Wallis’s test, followed by Dunn’s post-hoc test. Pearson’s correlation coefficient analysis was used for genotype-phenotype correlations.

## Results

### Determination of target-engaging dosages for NLX-112 and tandospirone administration in SCA3 mice

Despite solid descriptions of NLX-112 pharmacokinetics in rats (39), no dosing method had yet been established for chronic administration of this drug in mice. To determine two doses of the drugs that would allow sufficient exposure and 5-HT_1A_R engagement to carry on with the chronic studies, SCA3 mice received a single oral administration of different doses of NLX-112 (0.625, 2.5, and 10 mg/kg) and tandospirone (reference comparator, 2.5, 10 and 40 mg/kg) by OG **(Experimental design in Figure 1A**). Using LC-MS/MS, the drug concentration was measured in the plasma and brains of vehicle, NLX-112- and tandospirone-treated mice. These drugs were detected in plasma and brain samples in a dose-dependent manner (**Figure 1B and C, respectively**), with increased amounts of NLX-112 being detected in the periphery and central nervous system, compared to tandospirone. Importantly, the amounts of NLX-112 and tandospirone in the plasma of treated animals constituted a good readout for brain exposure.

**Figure 1.**
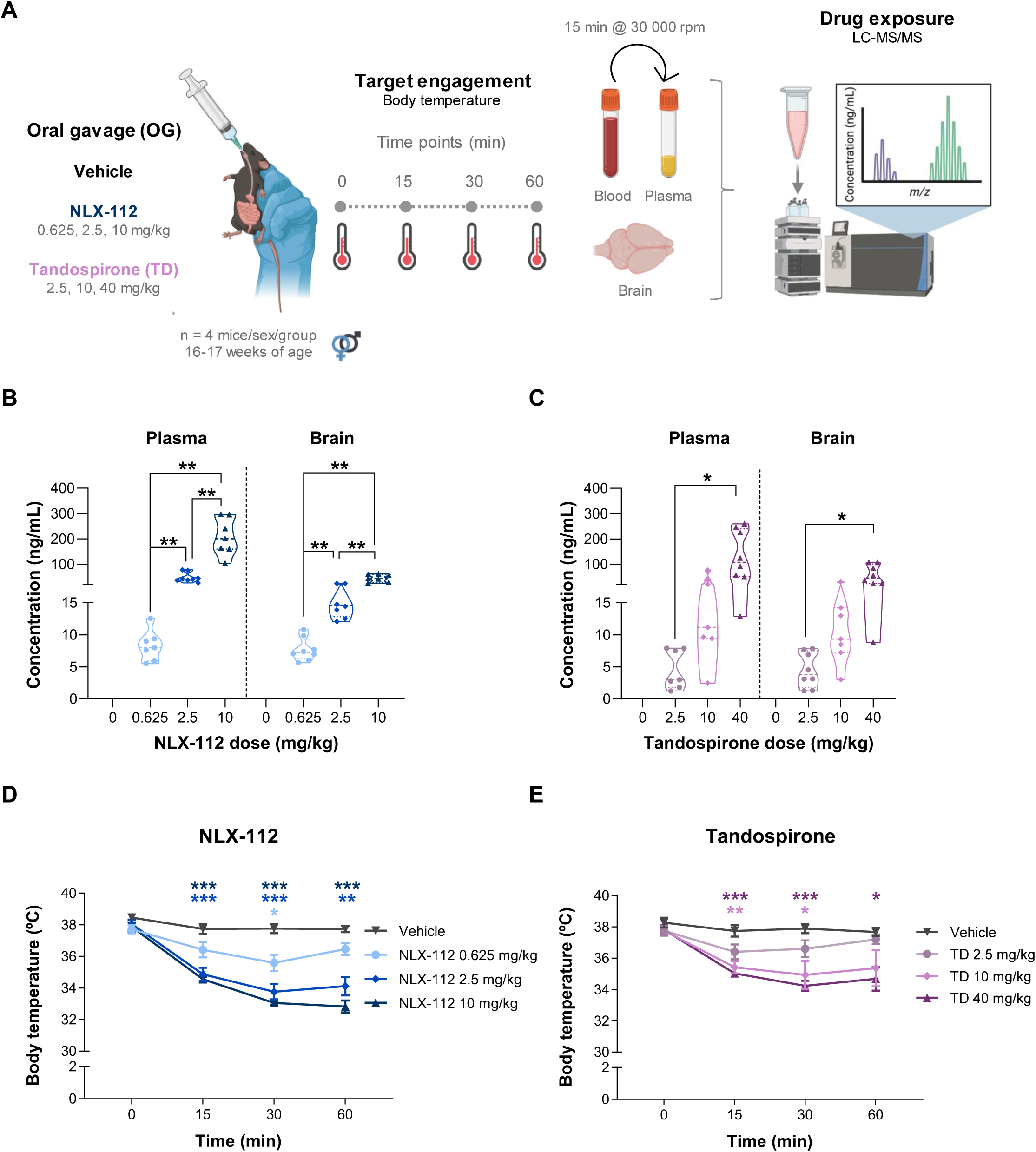
Drug exposure and validation of target engagement upon NLX-112 and tandospirone acute treatment. **(A)** Experimental design schematic representation (N = 70). SCA3 mice were given a single oral gavage of increasing doses of NLX-112 and tandospirone, and their core body temperature was measured at time 0, 15, 30 and 60 minutes after treatment. At minute 60, animals were euthanized, and the drug exposure was measured in plasma and brain samples by liquid chromatography-tandem mass spectrometry(LC-MS/MS).A dose-dependente detection of NLX-112 **(B)** and tandospirone **(C)** was achieved in the plasma and in the brain. Both NLX-112 **(D)** and tandospirone **(E)** showed dose-dependente target engagement by inducing hypothermia in mice. Two-way ANOVA or mixed-design ANOVA, Tukey, or Sidak post-hoc tests were computed. All data are expressed as group mean ± SEM (\**p*<0.05, \*\**p*<0.01, \*\*\**p*<0.001).

Next, drug-induced transient hypothermia upon acute administration, known to occur in rodents and mediated specifically by presynaptic 5-HT_1A_R in mice (40–43), was used as a biomarker for 5-HT_1A_R activation (43–49). This response is also elicited by 5-HT_1A_R agonists in humans (43). The rectal colonic temperature was recorded at 0, 15, 30, and 60 min upon drug administration. NLX-112 administration induced a decrease in body temperature after 15 min at higher doses (2.5 and 10 mg/kg) and after 30 min at the lower (0.625 mg/kg), concentration at which the temperature of the animals started to rise again 60 min after treatment (**Figure 1D**). The same trend was observed upon treatment with NLX-112 2.5 mg/kg, but not with the administration of 10 mg/kg, a dose at which the decreased body temperature remained low until the last measurement. The substantial decrease in mouse body temperature elicited by NLX-112 treatment at 10 mg/kg, challenged the principles of animal welfare, therefore this high dose was not used for further experiments.

Regarding tandospirone treatment, the higher doses (10 and 40 mg/kg) evoked a significant decrease in the animals’ body temperature, although not different among them. Upon tandospirone administration at 2.5 mg/kg, although not statistically significant, there was already a transient drug-induced decrease in the animals’ body temperature. All doses showed a trend towards body temperature recovery 60 min after treatment initiation (**Figure 1E**). Overall, the doses selected for NLX-112 and tandospirone induced significant decreases in body temperature, suggesting target engagement and 5-HT_1A_R activation.

### Tolerability of chronic NLX-112 and tandospirone administration in the drinking water for mice

After selecting the NLX-112 and tandospirone doses able to elicit acute 5HT_1A_R activation, the safety profile of these drugs was evaluated when administered chronically in the DW. SCA3 mice were treated for 28 days with NLX-112 (0.625 and 5 mg/kg) or tandospirone (2.5 and 20 mg/kg) and their welfare was monitored regularly (**Experimental design on Supplementary Figure 1A**). Animals’ body weight and water intake were measured weekly, with no signs of toxicity observed (**Supplementary Figure 1B and C, respectively**). Notably, no difference in water intake was observed in NLX-112- or tandospirone-treated mice compared to their vehicle-treated counterparts, suggesting the absence of an aversive taste or smell when these drugs are prepared in water. Observational welfare parameters, including natural grooming and nesting behaviors, dehydration, and fur appearance, were monitored during the entire experiment, with no signs of toxicity (**Supplementary Figure 1D**). Drug exposure analysis showed the presence of NLX-112 in the plasma and brain when administered in DW, in a dose-dependent manner; contrarily, tandospirone was not detected in brain samples at both doses tested, while only the highest dose was detected in plasma (**Supplementary Figure 1E**). Therefore, a dose quadruplication was applied for tandospirone (80 mg/kg), and the toxicity of this higher dose was also assessed in WT and SCA3 mice before proceeding to drug efficacy studies. No signs of toxicity were found for either genotype, as the animals showed no drug-induced decrease in body weight or other changes in welfare parameters upon two weeks of treatment (**Supplementary Figure 1F and D, respectively**). Regarding water intake, although there was a reduction in the water consumed by WT animals upon tandospirone water supplementation, the amount consumed by SCA3 animals was similar to that of WT animals exposed to the lower tandospirone concentrations (**Supplementary Figure 1G and C, respectively**).

### NLX-112 chronic treatment improved the balance of SCA3 mice

Considering the previous results, we advanced to efficacy studies using a low (LD) and high dose (HD) for each drug, administered *ad libitum* in the DW for 35 weeks: 0.625 and 5 mg/kg for NLX-112; and 20 and 80 mg/kg for tandospirone. Treatment started at 6 and ended at 41 weeks, and several motor behavior testing paradigms were applied, with an established periodicity according to the described phenotype for this SCA3 mouse model [(34); **Figure 2A**]. Because CAG repeat length in the *ATXN3* gene influences disease severity and varies among individuals (7, 37), we ensured group homogeneity. Importantly, in this study, no significant differences in CAG repeat length and range were found between vehicle- and NLX-112-treated SCA3 mice (**Figure 2B**), excluding such a confounding effect. Detection of LD and HD of NLX-112 in plasma and brain samples in a dose-dependent manner was confirmed after 35 weeks of treatment, assuring continuous drug exposure during the entire preclinical trial (**Figure 2C**). During the 35 weeks of treatment, NLX-112 administration via DW had no major impact on animals’ body weight gain (**Supplementary Figure 2A-B**), further confirming NLX-112 safety even when administered for a prolonged time. SCA3 animals showed a significant decrease in water intake compared to WT littermate animals (more pronounced in males), with no impact on water intake among SCA3 vehicle- and NLX-112-treated animals (**Supplementary Figure 2C-D**). Evaluation of core body temperature revealed a significant decrease in SCA3 mice, compared to their WT littermates, with no changes caused by prolonged administration of NLX-112 in the DW (**Supplementary Figure 2E-F**), regardless of the 5-HT_1A_R-mediated hypothermia seen upon acute exposure by OG (**Figure 1D**).

**Figure 2.**
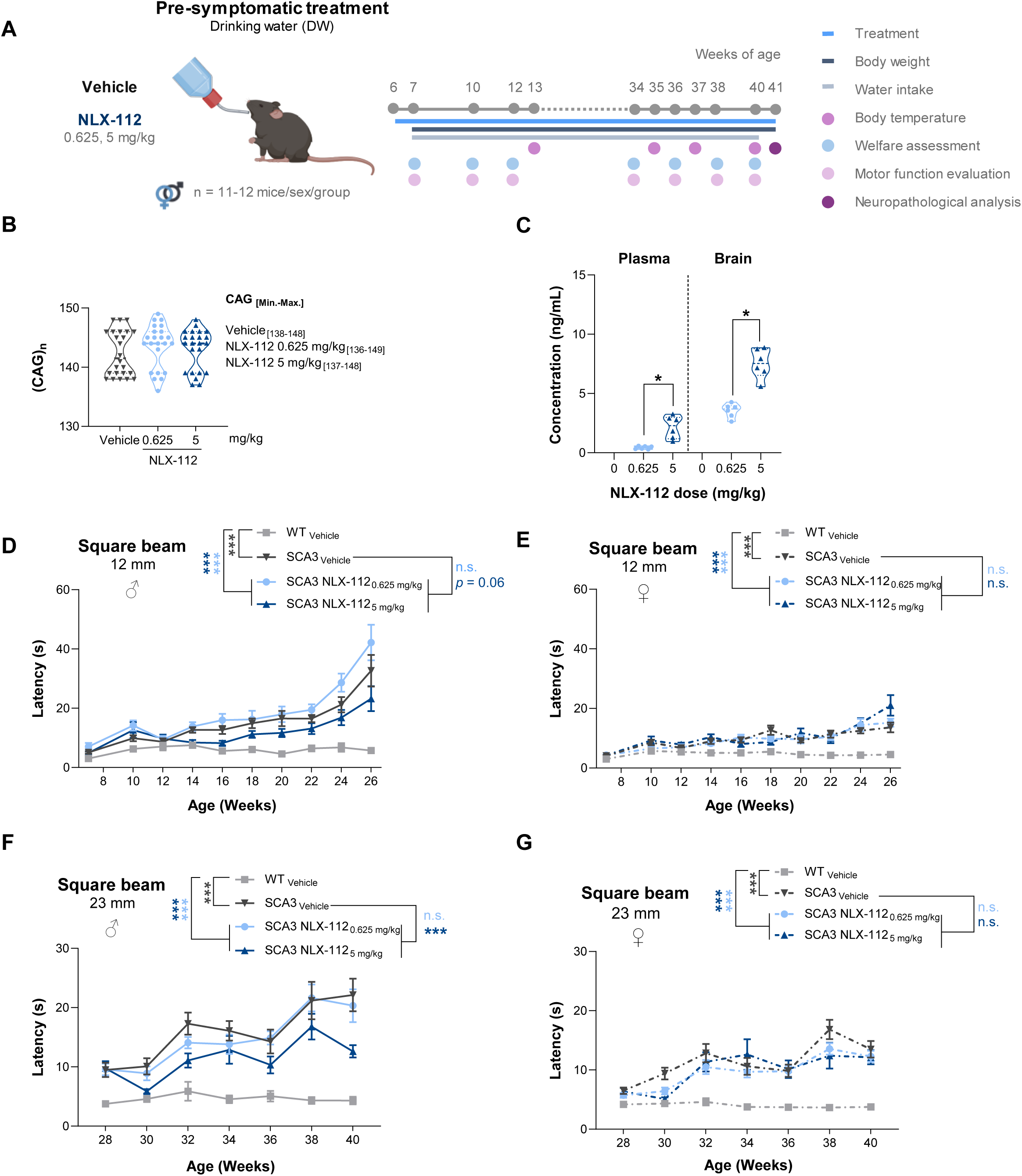
NLX-112 treatment improved SCA3 mice balance and coordination. **(A)** Schematic representation of the experimental design (N = 95). **(B)** The experimental groups showed similar CAG repeat length intervals.NLX-112 was detected in a dose-dependente manner in plasma and brain samples upon 41 weeks of exposure through the drinking water **(C)**. NLX-112 high dose administration to SCA3 mice showed a sex-dependente efficacy, by improving the latency to cross the 12 mm squared beam in males **(D)**, but not in females **(E).** The same result was found in a wider squared beam, where HD-NLX-112 treated male SCA3 mice presented a better performance **(F)**, when compared to vehicle-treated SCA3 mice, while the same was not observed for female mice **(G)**. Kruskal-Wallis test,Dunn’s post-hoc test for (B). Two-way ANOVA or mixed-design ANOVA, Tukey, or Sidak post-hoc tests for(C-G). All data are expressed as group mean ± SEM (\**p*<0.05, \*\*\**p*<0.001); n.s.–not significant).

Importantly, the administration of the HD of NLX-112 improved coordination and balance of SCA3 male mice in the 12 and 23 mm-squared and 20 mm-round beams (**Figure 2D and F and Supplementary Figure 2G, respectively**); whereas no therapeutic effect by NLX-112 was observed in SCA3 females in either dose or beam tested (**Figure 2E and G and Supplementary Figure 2H**). In contrast, administration of tandospirone failed to evoke any alteration in the balance of SCA3 mice. This lack of response occurred despite the absence of any observable adverse effects of the drug (**Figure 3 and Supplementary Figure 3**). Again, tandospirone was only detected at the periphery at both dosages, with tandospirone exposure in the brain samples being below the detection limit, mirroring the earlier findings regarding twenty-eight days of drug exposure in the DW (**Supplementary Figure 1D**).

**Figure 3.**
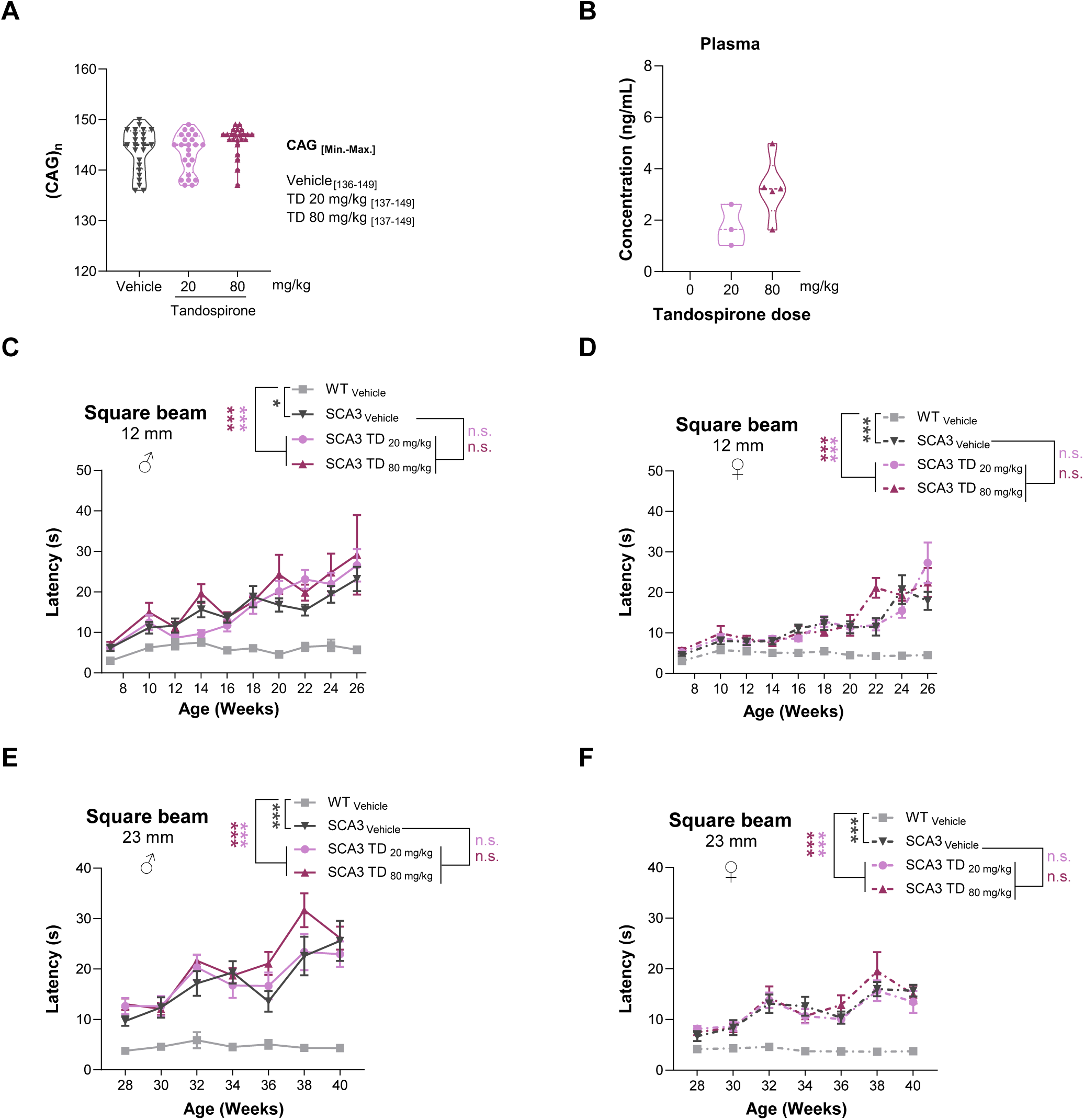
Tandospirone had no therapeutic Effect on SCA3 mice motor function. The CAG repeat length for all the experimental groups showed to be similar (N = 96) **(A)**. Both doses of tandospirone were detected in plasma samples **(B)**. The performance of SCA3 males and females treated with tandospirone on the 12 mm **(C-D)** and 23 mm **(E-F)** squared beams was close to SCA3-vehicle animals, demonstrating no therapeutic effect by tandospirone on SCA3 balance. Two-way ANOVA or mixed-design ANOVA, Tukey, Dunnet T3 or Sidak post-hoc tests were computed. All data are expressed as group mean ± SEM (\**p*<0.05, \*\**p*<0.01, \*\*\**p*<0.001, n.s. –not significant).

### NLX-112 improved neuropathological features in SCA3 mice

The analysis of brain tissue of SCA3 animals showed that NLX-112 treatment preserved the number of TH-positive (dopaminergic) neurons in the SN, which were similar to WT (**Figures 4A – upper panel, and B)**. NLX-112 treatment also mitigated astrocyte reactivity, decreasing glial fibrillary acidic protein (GFAP) staining intensity in the same brain region (**Figures 4A – lower panel and C**). This marked improvement in the SN neuropathology was independent of any impact on mutant ATXN3 nuclear inclusions, as end-stage aggregates are typically not detectable in the SN of this SCA3 model (34). However, NLX-112-treatment was able to modulate mutant ATXN3 aggregation in a region-dependent manner, as it increased the number of intranuclear inclusions in the DCN and it did not change their numbers in the PN nor in the SC (**Figure 4D**).

**Figure 4.**
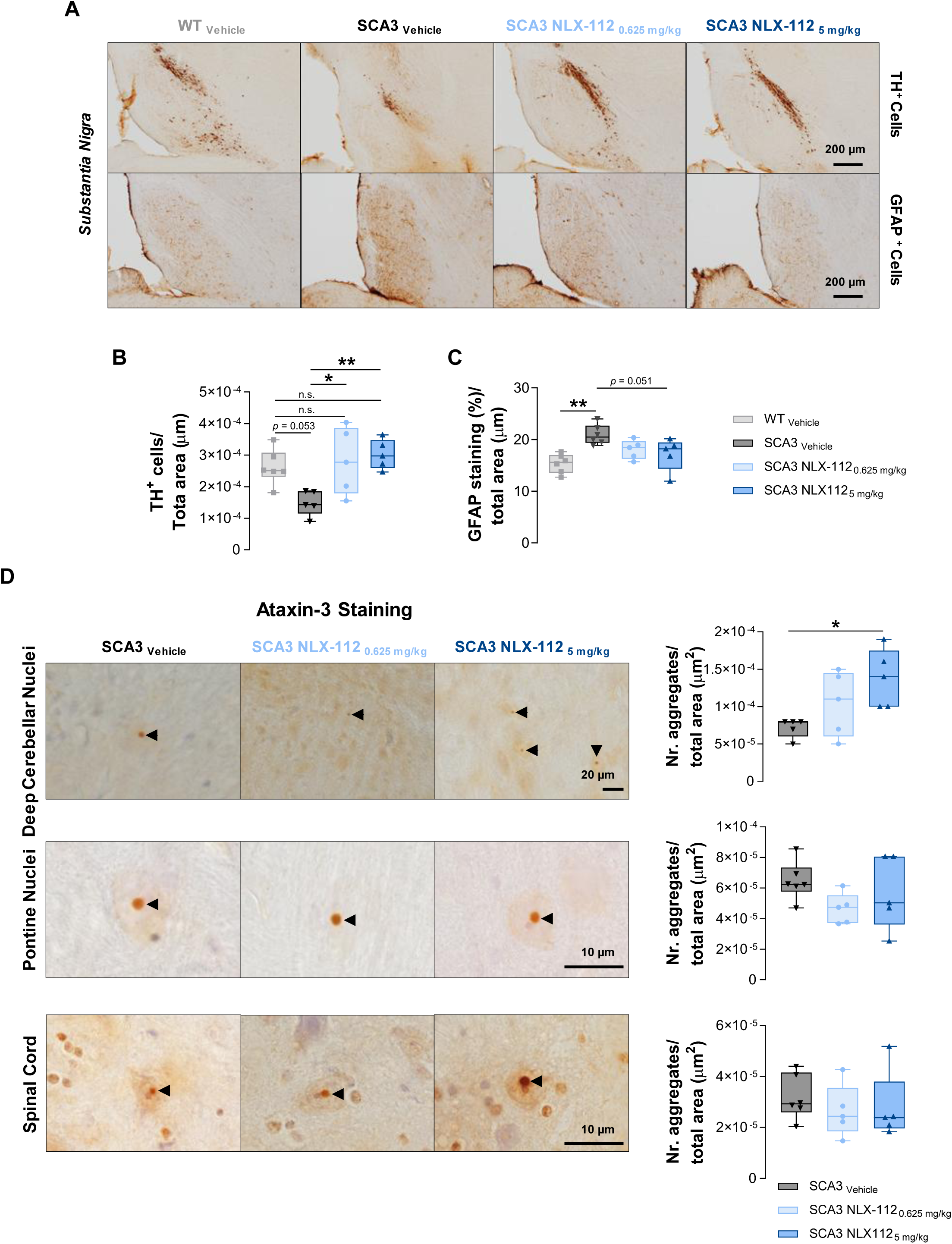
NLX-112 treatment is neuroprotective and has na impact in the number of ATXN3 nuclear inclusions, in a brain region-specific manner. Representative images for the antibody-specific immunostainings of TH^+^ and GFAP^+^ cells in the *substantia nigra* (SN) **(A)**. Both doses of NLX-112 were able to restore dopaminergic cells in the SN of SCA3 mice **(B)**. The high dose of NLX-112 decreased astrocyte reactivity in the SN of SCA3 mice **(C)**. Representative images for the antibody-specific ataxin-3 (ATXN3) staining in the deep cerebelar nuclei (DCN), pontine nuclei (PN) and spinal cord (SC) **(D)**. SCA3 mice treated with the high dose of NLX-112 showed increased number of nuclear ATXN3 aggregates in the DCN, but not in the PN or SC **(D).** n = 5-6 animals/group, 5-10 slices/animal/experimental group, mixed sexes. Arrowheads: ATXN3 nuclear inclusions. Scale bar 200 μm for (A), 20μm (D – first frame) and 10 μm (D – second and third frames). One-way ANOVA, Tukey post-hoc test. All data are expressed as group mean ± SEM (\**p*<0.05, \*\**p*<0.01).

## Discussion

The search for novel treatments for spinocerebellar ataxias (SCAs) represents a pressing unmet medical need, as there are currently no effective disease-modifying or symptomatic therapies for this group of disorders. The present study, focusing on NLX-112 and tandospirone, suggests that selective targeting of 5-HT_1A_Rs may represent a promising strategy for improving symptoms and slowing disease progression in individuals living with SCA3.

The core findings of the study are as follows: (i) Upon chronic administration in the DW for 35 weeks, starting prior to the onset of ataxia-related motor deficits, NLX-112, at the tested doses, improved motor coordination in male (but not female) mice. Tandospirone did not have significant effects. (ii) NLX-112 attenuated various measures of cellular neuropathology in SCA3 mice. Specifically, it reduced dopamine neuron loss and astrocyte reactivity in the SN, which are important markers of neuroprotection in the central nervous system (CNS). NLX-112 treatment also increased the number of nuclear mutant ATXN3 end-stage aggregates – likely reducing their neurotoxicity – in a brain region-specific manner. (iii) Prolonged drug administration was safe and well-tolerated for SCA3 mice, mirroring observations in human trials (31, 50).

Targeting serotonergic signaling represents a promising therapeutic approach for the treatment of SCA3 and other forms of ataxia. For example, citalopram treatment attenuated SCA3 in a *C. elegans* model via the SERT/MOD-5 transporter and 5-HT_1A_/SER-4 receptor (14). Citalopram was also active in two mouse models of SCA3 (14, 15, 37), showing beneficial effects on the animals’ mobility/coordination, as well as on the aggregation of mutant ATXN3. These observations therefore provided a robust rationale to explore the effects of targeting 5-HT_1A_R with the highly selective agonist, NLX-112 and the partial agonist, tandospirone. Initial experiments confirmed that NLX-112 and tandospirone engaged the 5-HT_1A_R in the SCA3 mice, as indicated by decreased body temperature, and the compounds were well-tolerated when administered chronically, showing no adverse effects at the selected doses. NLX-112 and tandospirone were then tested in chronic administration protocols, in which treatment was started before the onset of CNS-related core motor deficits. This is because, as the disease progresses, the capacity of some drug treatments to prevent aggregation and neurodegeneration may be compromised, as suggested by a previous study in *C. elegans* model of SCA3: NLX-112 ameliorated motor function and decreased mutant ATXN3 aggregation after both acute and chronic treatment, but tandospirone failed to do the same (33). In spite of ongoing trend that SCA3 patients are usually diagnosed after the first symptoms appear and only treated subsequently, there is an opportunity for genetic testing prior to disease onset (51). Based on this and considering that, as for other diseases “Time is Brain”, it is crucial to understand the potential therapeutic effects of novel drugs under conditions where treatment initiation occurs when disease symptoms are not yet present or at early symptomatic stages.

In the present study, administration of NLX-112 resulted in significant improvements in motor coordination and balance in male mice. These beneficial effects were more pronounced during the later stages of the disease, with NLX-112 treatment attenuating symptom severity, as evidenced by a reduced latency to cross the beam from 28 weeks onward. It is unclear why the female mice did not show similar improvement in motor function as the males, but this sex-dependent effect of NLX-112 on motor function was possibly associated with differences in the sensitivity of serotonergic systems between males and females or SCA-specific changes in 5-HT_1A_R regulation. We hypothesize that this could be due to early serotonergic events during brain development, such as differences in the density or affinity of 5-HT_1A_ auto- and hetero-receptors related to sex and/or the influence of gonadal hormones in both sexes, as indicated by previous studies (52, 53). Distinct responses to 5-HT_1A_R agonists and differential gene expression in 5-HT neurons were also reported among sexes and the estrous cycle (54–63). Studies in humans also revealed region-specific alterations in 5-HT_1A_R binding changes with aging, associated with sex-specific receptor density and affinity (64), and that the rate of 5-HT synthesis is higher in healthy male humans than in females (65). Based on this, it may be speculated that higher doses of NLX-112 would have been effective in the female mice. However, more research is required to clarify the sex differences in the serotonergic system, which could have implications for the use of serotonergic drugs.

In contrast to NLX-112, chronic tandospirone treatment failed to improve the motor function of SCA3 mice. This may be due to the low exposure levels of tandospirone in the brain, likely arising from its short metabolic half-life, which might require more frequent pulsatile dosing. It is also likely that the lack of activity of tandospirone may be due to its partial agonist properties at 5-HT_1A_Rs, which may be insufficient to elicit optimal therapeutic-like benefit in the mice.

Chronic administration of NLX-112 restored several disease-specific measures of cellular neuropathology in SCA3 mice. Specifically, NLX-112 reduced dopamine (TH-positive) neuron loss and astrocyte reactivity in the SN, which are important markers of neuroprotection in the CNS. The neuroprotective effect of NLX-112 is supported by data from other 5-HT_1A_R agonists, for example 8-OH-DPAT, which induced astrocyte proliferation and increased antioxidative molecules, as well as reversed the dopaminergic neurodegeneration *in vitro* and *in vivo* experiments in Parkinsonian mouse models (66, 67). Intriguingly. NLX-112 also increased the number of nuclear mutant ATXN3 end-stage aggregates in the DCN, suggesting that the most relevant mutant ATXN3 toxic species might be more soluble and/or smaller aggregates than the ones that are microscopically visible (resolution limit of ∼0.2 mm) and that were quantified here. No impact on end-stage mutant ATXN3 protein aggregates was noted in the PN and SC neurons.

This region-specific effect of NLX-112 administration might be due to the distinct 5-HT_1A_ receptor expression, co-expression pattern and/or activation of this and other 5-HT receptors in different CNS regions. This regulates serotonergic neurons’ excitability, as was previously well documented for the 5-HT_1A_ and 5-HT_2B_ receptors co-clustering (68); decoding this is essential to understand the underlying effects of NLX-112 in the context of SCA3 (69–71).

Overall, these results reinforce the assertion that selective targeting of 5-HT_1A_R could be a useful therapeutic approach for SCA3 and other movement disorders (14, 15, 20, 21, 37, 72–75). NLX-112 also exhibits antiparkinsonian (76) and antidyskinetic properties (26, 76–78) in animal models of PD and significantly reduced both dyskinesia and parkinsonism in a proof-of-concept clinical trial (32, 79). This highlights the potential benefits of developing 5-HT_1A_R full agonists, such as NLX-112, for the treatment of movement disorders and supports its translatability from animal studies to clinical trials in SCA3. It should be noted that NLX-112 has been granted Orphan Drug designation, in both the European Union and in the United States, as a treatment for Spinocerebellar Ataxia.

## Acknowledgments

We are grateful to current members of the Translational Neurogenetics team for their critical analysis of the data and helpful discussions.

## Authors’ Roles

Bruna Ferreira-Lomba: execution, writing.

Sara Guerreiro: execution, writing.

Sara Duarte-Silva: design, execution, analysis, writing.

Daniela Cunha-Garcia: execution, analysis, writing.

Stéphanie Oliveira: execution.

Cármen Vieira: execution.

Joana Pereira-Sousa: execution.

Daniela Vilasboas-Campos: execution.

André Vidinha-Mira: execution.

Daniela Monteiro-Fernandes: execution.

Mark A Varney: design, funding acquisition.

Mark S Kleven: design, management, supervision, editing of final version of the manuscript.

Adrian Newman-Tancredi: design, management, supervision, editing of final version of the manuscript, funding acquisition.

Andreia Teixeira-Castro: execution, design, management, supervision, editing of final version of the manuscript.

Patrícia Maciel: design, management, supervision, editing of final version of the manuscript, funding acquisition.

## Financial Disclosures of all authors

Mark A. Varney, Mark S. Kleven and Adrian Newman-Tancredi are employees and/or stockholders of Neurolixis Inc. Patrícia Maciel and Andreia Teixeira-Castro possess the intellectual property rights of the patent entitled “Citalopram or escitalopram for use in the treatment of neurodegenerative diseases” (Patent Number: 242.8, Portugal; US11033526B2, USA). The other authors have no relevant disclosures.

This work was funded by: the Department of Defense (DoD) - U.S. Army Medical Research and Materiel Command (Award Number: W81XWH-19-1-0638) and by Portuguese funds, through the Foundation for Science and Technology (FCT), under projects UID/06304/2023 and LA/P/0050/2020 (DOI 10.54499/LA/P/0050/2020) and by the project NORTE2030-FEDER01786400, supported by Norte Portugal Regional Operational Programme (NORTE 2030), under the PORTUGAL 2030 Partnership Agreement, through the European Regional Development Fund (ERDF) and by ICVS Scientific Microscopy Platform, member of the national infrastructure Portuguese Platform of Bioimaging (PPBI) (PPBI-POCI-01-0145-FEDER-022122). Additionally, this study was supported by the funding received by Andreia Teixeira-Castro from FCT Exploratory Project (2023.15102.PEX), Early Career Investigator Grant by National Ataxia Foundation (NAF) and Ataxia UK and by the following individual fellowships provided by FCT: Scientific Employment Stimulus (CEEC) — Individual Call position to Sara Duarte Silva (CEECIND/00685/2020); PhD fellowships of Bruna Ferreira-Lomba (2024.01049.BD), Sara Guerreiro (2022.11724.BD), Daniela Cunha-Garcia (2021.08121.BD), Cármen Vieira (2022.11176.BD), Daniela Vilasboas-Campos (SFRH/BD/147826/2019) and Daniela Monteiro-Fernandes (SFRH/BD/147947/2019).

---

**Supplementary Figure 1.**
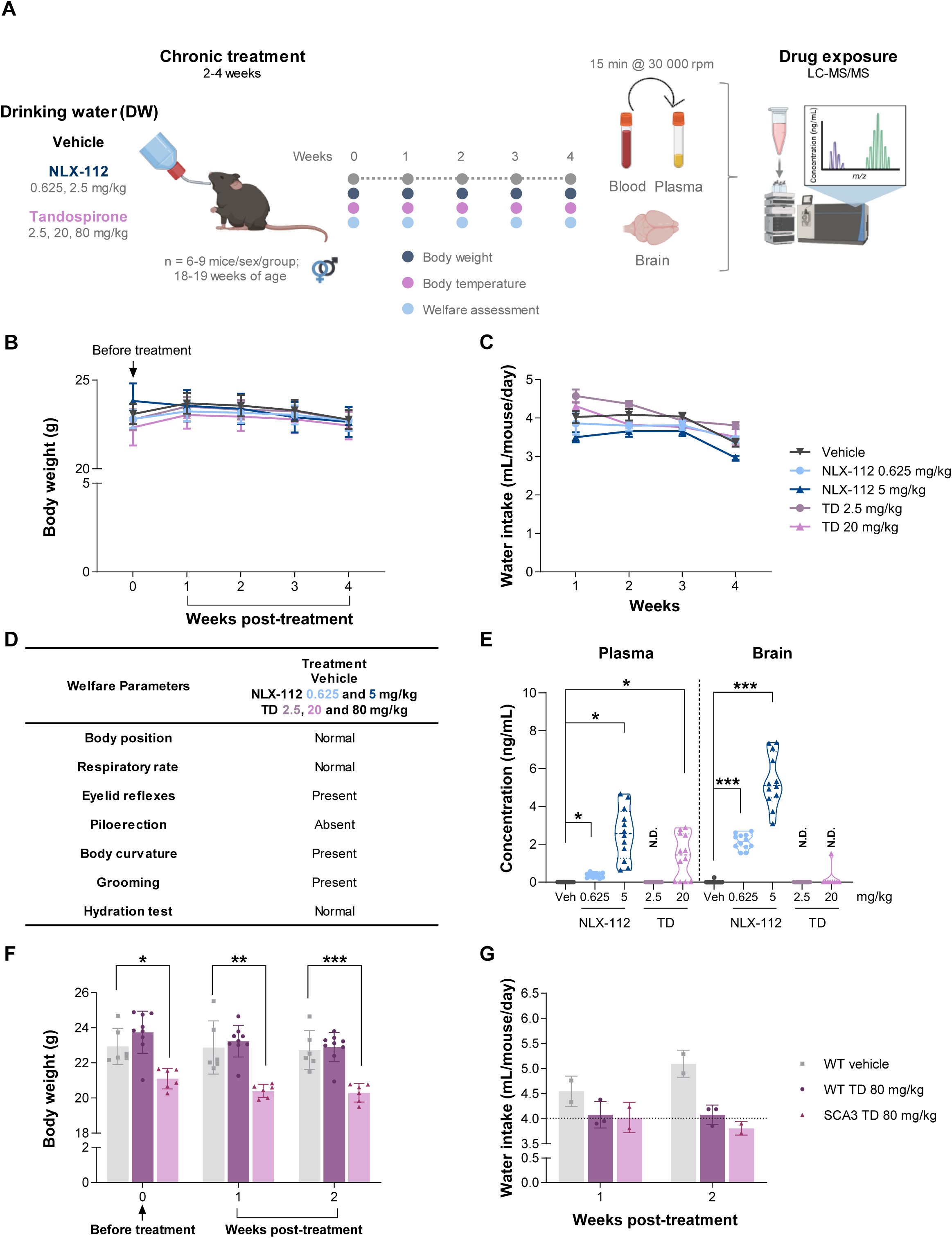
NLX-112 and tandospirone administration over a 28-day period by the drinking water was safe to mice. **(A)** Schematic representation of the experimental design of the sub-chronic NLX-112 andt andospirone treatment through the drinking water (N = 60, n = 6 mice/sex/group for **B**, **C** and **E**; N = 21, n = 6-9 mice females/group for **D**, **F** and **G**). These drugs caused no alterations in animals’ body weight **(B)** or water intake **(C).** A panel of welfare tests was performed and all mice demonstrated the expected behavior **(D).** Four weeks post-treatment, plasma and brain samples were collected to measure the levels of the drugs **(E)**. Dose-dependente NLX-112 was detected in both sample types, but only the high dose of tandospirone was detected in the plasma; none of the doses was detected in the brain. A dose quadruplication was applied, and 80 mg/kg of tandospirone was first tested for safety. A 2-weeks treatment to both WT and SCA3 mice showed no signs of toxicity given by their normal body weight **(F)** and water intake **(G).** Two-way ANOVA or mixed-design ANOVA,Tukey, or Sidak post-hoc tests were computed. All data are expressed as group mean ± SEM (\**p*<0.05, \*\**p*<0.01, \*\*\**p*<0.001). N.D. – not detected.

**Supplementary Figure 2.**
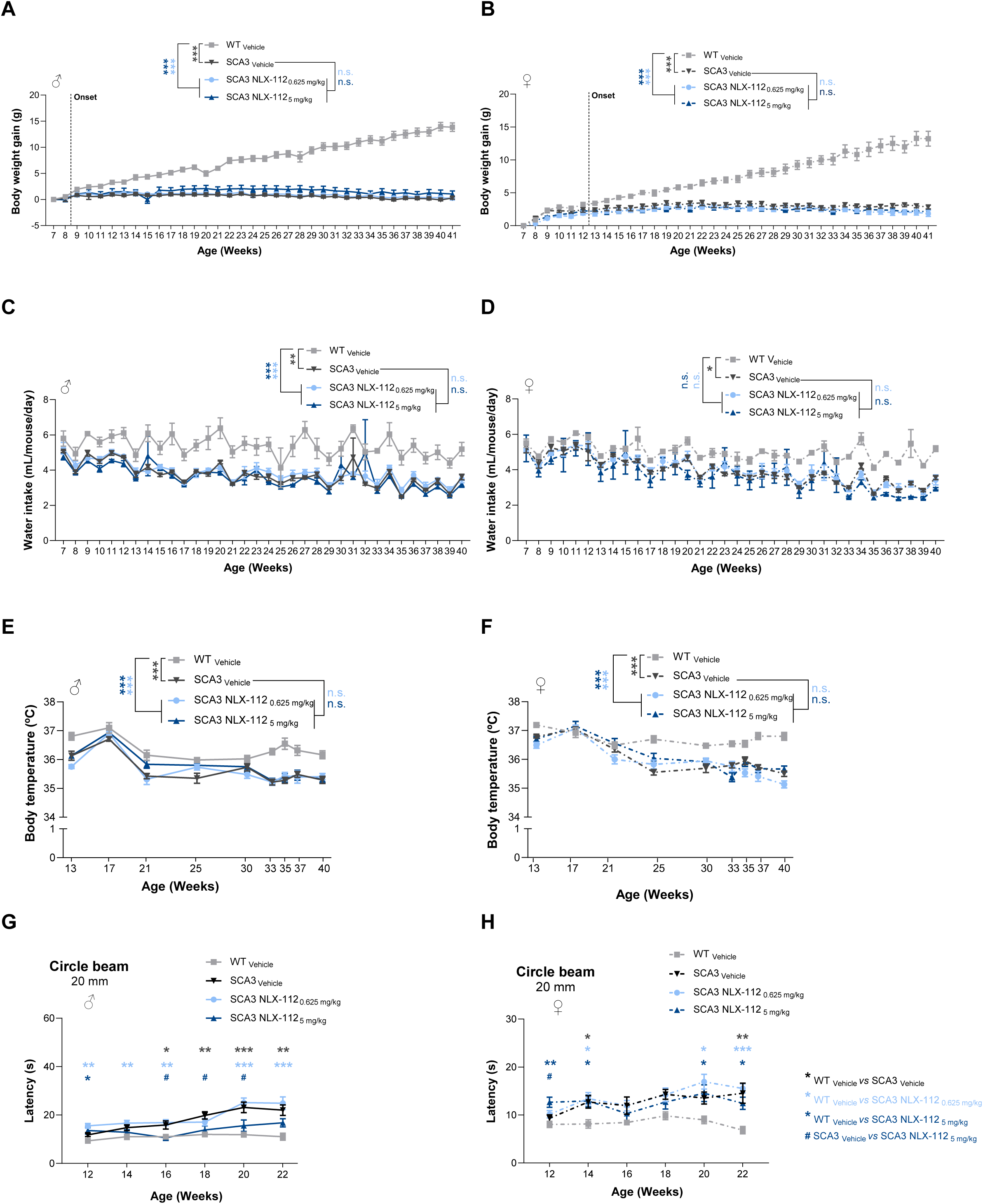
NLX-112 chronic treatment is safe and well-tolerated. Two doses of NLX-112 were administered through the drinking water over a period of 35 weeks to SCA3 mice (N = 95). No signs of toxicity were observed during the entire preclinical trial, and for both males and females no changes were found in their body weight gain **(A-B)**, water intake **(C-D)** and core body temperature **(E-F)**.Higher dose of NLX-112 improved the motor performance of male mice on the 20 mm circle beam in specific time points but had no impact on females **(G-H).** Two-way ANOVA or mixed-design ANOVA, Tukey, or Sidak post-hoc tests were computed. All data are expressed as group mean ± SEM (\**p*<0.05, \*\**p*<0.01, \*\*\**p*<0.001, n.s.– not significant).

**Supplementary Figure 3.**
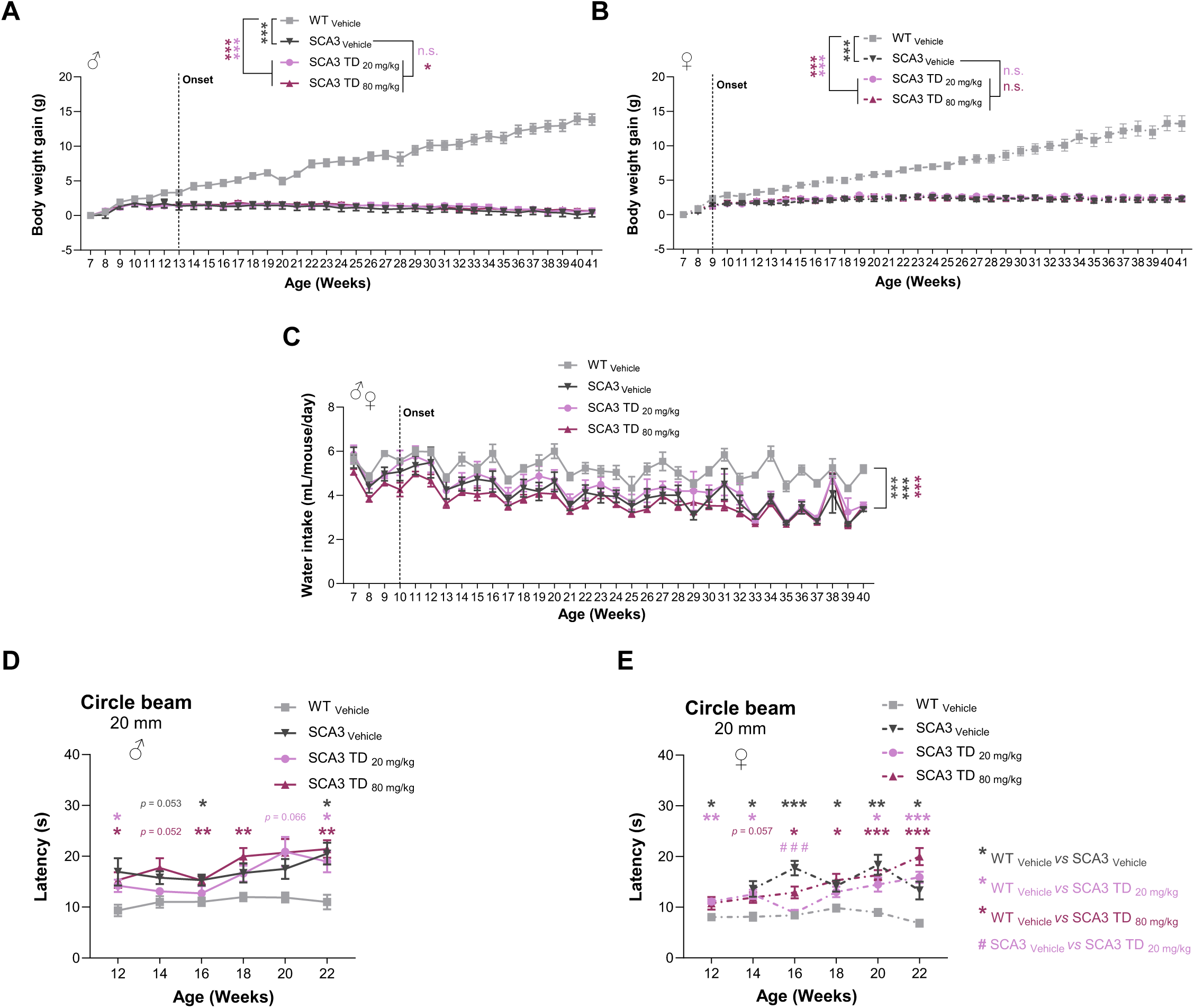
Tandospirone treatment generally failed to produce any Effect on SCA3 mice. Tandospirone chronic treatment had na impact on the body weight gain of male SCA3 mice at the higher dose tested (**A**), but not in females, where the rate of body weight gain is comparable to SCA3 vehicle mice (**B**). Tandospirone had no impact on water intake of SCA3 mice (**C**). The performance of SCA3 males and females treated with tandospirone on the 20 mm circle beam was like SCA3-vehicle animals (**D-E**). Two-way ANOVA or mixed-design ANOVA, Tukey, Dunnet T3 or Sidak post-hoc tests were computed. All data are expressed as group mean ± SEM (\**p*<0.05, \*\**p*<0.01, \*\*\**p*<0.001, n.s. – not significant).

## Notes

### Competing Interest Statement

Mark A. Varney, Mark S. Kleven and Adrian Newman-Tancredi are employees and/or stockholders of Neurolixis Inc. Patricia Maciel and Andreia Teixeira-Castro possess the intellectual property rights of the patent entitled -Citalopram or escitalopram for use in the treatment of neurodegenerative diseases- (Patent Number: 242.8, Portugal;
US11033526B2, USA). The other authors have no relevant disclosures.

