## Supplementary methods for "Efficacy of chronic 5-HT_1A_ receptor agonism by NLX-112 in a mouse model of Spinocerebellar Ataxia type 3"

**Running title**: NLX-112 treatment to mitigate SCA3

Ferreira-Lomba B[1,2*], MSc; Guerreiro S[1,2*], MSc; Duarte-Silva S[1,2*], PhD; Cunha-Garcia D[1,2], MSc; Oliveira S[1,2], PhD; Vieira C[1,2], MSc; Pereira-Sousa J[1,2], PhD; Vilasboas-Campos D[1,2], MSc; Vidinha-Mira A[1,2], MSc; Monteiro-Fernandes D[1,2], MSc; Varney MA[3], PhD; Kleven MS[3], PhD; Newman-Tancredi A[4$], PhD/DSc; Teixeira-Castro A[1,2$], PhD; Maciel P[1,2$#], PhD.

[1] Life and Health Sciences Research Institute (ICVS), School of Medicine, University of Minho, Braga, Portugal.
[2] ICVS/3B's – PT Government Associate Laboratory, Braga/Guimarães, Portugal.
[3] Neurolixis Inc., Park Ridge, NJ 07656, USA.
[4] Neurolixis SAS, 81100 Castres, France.

* Equal contribution
[$] Co-senior authorship
[#] Corresponding author

### Housing and Health Conditions

The animals had a Specified Pathogen Free health status and were maintained under standard conditions in the vivarium, namely: artificial 12 h light/dark cycle (lighting from 8 AM to 8 PM), 21 ± 1°C and relative humidity of 50-60 %. A maximum of five mice were housed by genotype and treatment conditions in filter-topped polysulfone cages (Tecniplast), with corncob bedding (Scobis Due, Mucedola S.r.l.), enriched with soft tissues and shredded paper for nesting behavior. The cages were placed on a standard rack (Tecniplast). The animals were fed with a standardized diet, during gestation, postnatally (4RF25, Mucedola S.r.l.) and after weaning at 3 weeks-old or after arriving at the vivarium (4RF21, Mucedola S.r.l.). The animals ate and drank water *ad libitum*.

### NLX-112 and Tandospirone Levels Determination

To determine NLX-112 and tandospirone levels, the brains were mechanically homogenized in PBS. The plasma and brain samples were subjected to an extraction procedure: tolbutamide solution at 0.1 μg/mL or tandospirone-$d_8$ solution at 0.02 µg/mL (internal standards for NLX-112- or tandospirone-treated samples, respectively) or acetonitrile for S0 were added; the samples were mixed for 30 s, centrifuged at 20,000 g and 4 ºC for 5 min and the supernatant was transferred. Then, RP868 ($H_2O$/methanol; 75/25 % v/v) or acetonitrile was added (for NLX-112- or tandospirone samples, respectively). The samples were mixed for 5 min, centrifuged at 2,500 g and 4°C for 5 min and then injected into the LC-MS/MS. High performance liquid chromatography (HPLC) method was performed using: the ACQUITY UPLC™ BEH $C_{18}$ (2.1 x 50 mm, 1.7 µm; Waters Corporation, part number: 186002350) or Kromasil (50 x 3 mm, 5 µm) chromatographic column, a flow rate (mL/min) of 0.6 or 0.8 and the mobile phases were constituted by 1 mM ammonium acetate + 0.025 % formic acid in $H_2O$/acetonitrile (95/5 %) (phase A) and 1 mM ammonium acetate + 0.025 % formic acid in $H_2O$/acetonitrile (5/95 %) or $H_2O$ + 0.1% formic acid (phase A) and acetonitrile + 0.1% formic acid (phase B), for NLX-112- or tandospirone-treated samples, respectively. The LC-MS/MS conditions for NLX-112- or tandospirone-treated samples (respectively) were the following: detector – API5500 (Sciex)/8060 (Shimadzu); ionisation mode – ESI+; acquisition mode – MRM; mass transition – NLX-112: 394.2/374.2, tolbutamide: 271.1/155.0, tandospirone: 384.1/122.15 and tandospirone-$d_8$: 392.1/122.15. The lower quantification limit was <0.1 ng/mL or <0.3 ng/g for NLX-112 and <1 ng/mL or <3 ng/g for tandospirone.

**Beam Walk Test**

Polyvinyl chloride (PVC) beams with different formats (circle or square) and thicknesses (diameters) were used: 12 mm square beam (7-26 weeks of age, training beam) and 20 mm cylindrical beam (12-22 weeks of age). These beams were replaced by a 23 mm square beam (at 28-40 weeks of age, easier to traverse), when the SCA3 mice phenotype worsened and most of them were unable to complete the task without falling. The animals were trained over 3 consecutive days (2 trials/animal) and evaluated on the 4$^{th}$ day (2 trials/beam/animal and 2 fails allowed/beam/animal), counting the time needed to perform the test and removing the immobility time. The trial was invalid if the mouse fell or turned around during the test. The behavioral tests were done during the lighting period.

**Neuropathology Evaluation**

The ATXN3-positive intranuclear inclusions and TH-positive cells were quantified in several regions (deep cerebellar nuclei – DCN; pontine nuclei – PN; thoracic and lumbar spinal cord – SC; *substantia nigra* – SN) of treated- and non-treated-SCA3 mice (n= 5-6 animals, 5-10 slices/animal/experimental group, mixed sexes), using the Olympus BX51 stereological microscope with the Visiopharm Integrator System software. The contour of the region of interest was made using a 4x objective to draw the mask and to count the cells at 40x objective (1.0 numerical aperture oil immersion), in random counting frames of 80 µm x 80 µm in all the regions (fraction of sampling/slice: 100 % of DCN, PN, thoracic and lumbar SC and SN). Then, the total of inclusions or cells counted was divided by the area of the brain region ($\mu m^2$), giving the density of inclusions or cells per $\mu m^2$. For quantification of glial fibrillary acidic protein (GFAP)-positive cells intensity, mosaic pictures were obtained using the automatic Olympus cellSens Dimension Software (instant MIA, objective 10x) in the Olympus Upright BX61 microscope (0.16 numerical aperture objective), equipped with a DP74 camera. Using the Fiji software (ImageJ, version 1.545), the region of interest was drawn (ROI area) and a macro was used to apply a threshold (Otsu Dark algorithm), retrieving an automatic minimum value cut-off that selects “foreground”, eliminating the background of the image and allowing to specifically quantify the intensity of the marker in the chosen area. All experimenters performing the quantifications were blind to the experimental group.
